# CONSERVED LIFE HISTORY PHENOTYPES AMONG HISTORICALLY ISOLATED CLADES IN THE CHIRAL LIVEBEARER FISH OF LOWER CENTRAL AMERICA

**DOI:** 10.1101/2025.11.16.688739

**Authors:** E. Elias Johnson, Erik S. Johnson

**Affiliations:** Evolutionary Ecology Laboratories, Department of Biology and Bean Life Science Museum, Brigham Young University, Provo, UT, 84602, USA; Department of Biology and Whitney R. Harris World Ecology Center, University of Missouri–St. Louis, MO, 63121, USA

## Abstract

Lower Central America contains some of the highest freshwater fish diversity on the planet. Yet, species by species, we still know remarkably little about factors that have created this diversity and the threats that could put it at risk. The chiral livebearer (*Xenophallus umbratilis*) exemplifies these deficits. This freshwater livebearing fish species primarily occupies habitats along the high-elevation fringes of large volcanoes in the central cordillera of Costa Rica.

Previous work suggests that current populations have been isolated by repeated marine incursions beginning around 5 mya. Genetic data point to four putative clades that could be considered evolutionarily distinct. Unfortunately, beyond this we know little about adaptive evolution of demographically relevant traits among these clades, making it difficult to determine the conservation status of this species overall. Moreover, a recent IUCN assessment of this species was limited to only a portion of its geographic range and lacked population level information from many locations where it occurs. To address these gaps, here we describe the life history of this species, taken from 23 collections made across its geographic range from locations where this fish is locally abundant. This allowed us to determine if life history traits differ among populations coincident with known population structure. Despite long periods of isolation among clades, we found only modest evidence for divergence in life history phenotypes, and this was limited to just two of the four clades. These data suggest that life history traits are mostly conserved among populations, data that are critical to future conservation planning.

## Introduction

Few places on earth house greater freshwater fish diversity than the Neotropics of Lower Central America. This region, where the North and South American land masses connected, facilitating the Great American Biotic Interchange, is also a freshwater fish biodiversity hotspot (Bussing, 2002; Smith & Bermingham, 2005; Bagley & Johnson, 2014; Rodríguez *et al*., 2022; Fernando Gómez-Martínez *et al*., 2025). Several factors have likely contributed to the origin and maintenance of this diversity, including ecological interactions, such as competition and predation (Jennions & Telford, 2002; Johnson & Zuñiga-vega, 2009), as well as vicariant factors such as tectonic activity, volcanism, and isolation associated with marine incursion events (Smith & Bermingham, 2005; Jones & Johnson, 2009; Bagley *et al*., 2017). In fact, current measures might underestimate freshwater fish diversity in this region as species counts continue to increase due to ongoing taxonomic work (e.g., Matamoros *et al*., 2013; Angulo *et al*., 2018; Arroyave *et al*., 2024). Lower Central America is clearly a cradle for tropical fish diversification.

Unfortunately, this region is also at risk of losing freshwater fish biodiversity due to anthropogenic causes, even before this diversity is completely understood and described. A recent IUCN assessment of freshwater fishes in Lower Central America showed that up to 28% of described species were viewed as threatened with extinction, with dozens of additional species being data deficient so that we cannot adequately determine their conservation status (Rodríguez *et al*., 2022). Fishes in the family Poeciliidae, commonly known as the livebearers, are particularly vulnerable because they often occupy very specific niches in the tropical freshwater landscape, making them susceptible to the negative effects of human activity (Stockwell & Henkanaththegedara, 2011; Lyons, 2019). Deforestation, species invasions associated with aquaculture, mining, and pollution can all negatively impact these fishes (Stockwell & Henkanaththegedara, 2011; Mena et al. 2014; Lyons, 2019; Elias et al., 2022). One of our best approaches in conservation planning has been to study livebearing fishes using genetic, ecological, and evolutionary data to determine species conservation status (Rodríguez *et al*., 2022). Yet even the most basic biology for most taxa is still lacking. What is needed are studies that examine the evolutionary history of these fishes to determine how best to conserve standing diversity and how best to protect future opportunities for further diversification.

Here, we address the conservation status of the livebearing fish *Xenophallus umbratilis* (Figure 1). We demonstrate how phylogeographic data can be combined with life history data to better understand conserving diversity within Neotropical freshwater fishes. *Xenophallus umbratilis* is a monotypic genus (hereafter *Xenophallus*) with a relatively narrow distribution, most commonly around the high-elevation fringes of volcanoes in northwestern Costa Rica and southern Nicaragua (Bussing, 2002). Previous research shows that this species can be divided into four distinct evolutionary clades that appear to have become isolated during repeated marine incursion events associated with glacial melting cycles since at least the Pliocene (Jones & Johnson, 2009). Without additional information, a cautious approach might be to protect representatives from each of these clades as unique conservation units. Here, we ask if this conclusion is warranted based on life history traits within this species. Moreover, we ask if these clades of *Xenophallus*, some isolated as long as 4.5 million years ago, have evolved divergent life history strategies. We focus on life histories because these traits evolve rapidly, and ultimately, contribute to demographic processes that control the decline, rise, or persistence of populations (Heppell *et al*., 2000; Roff, 2002; Johnson & Zuñiga-Vega 2009).

**Figure 1.**
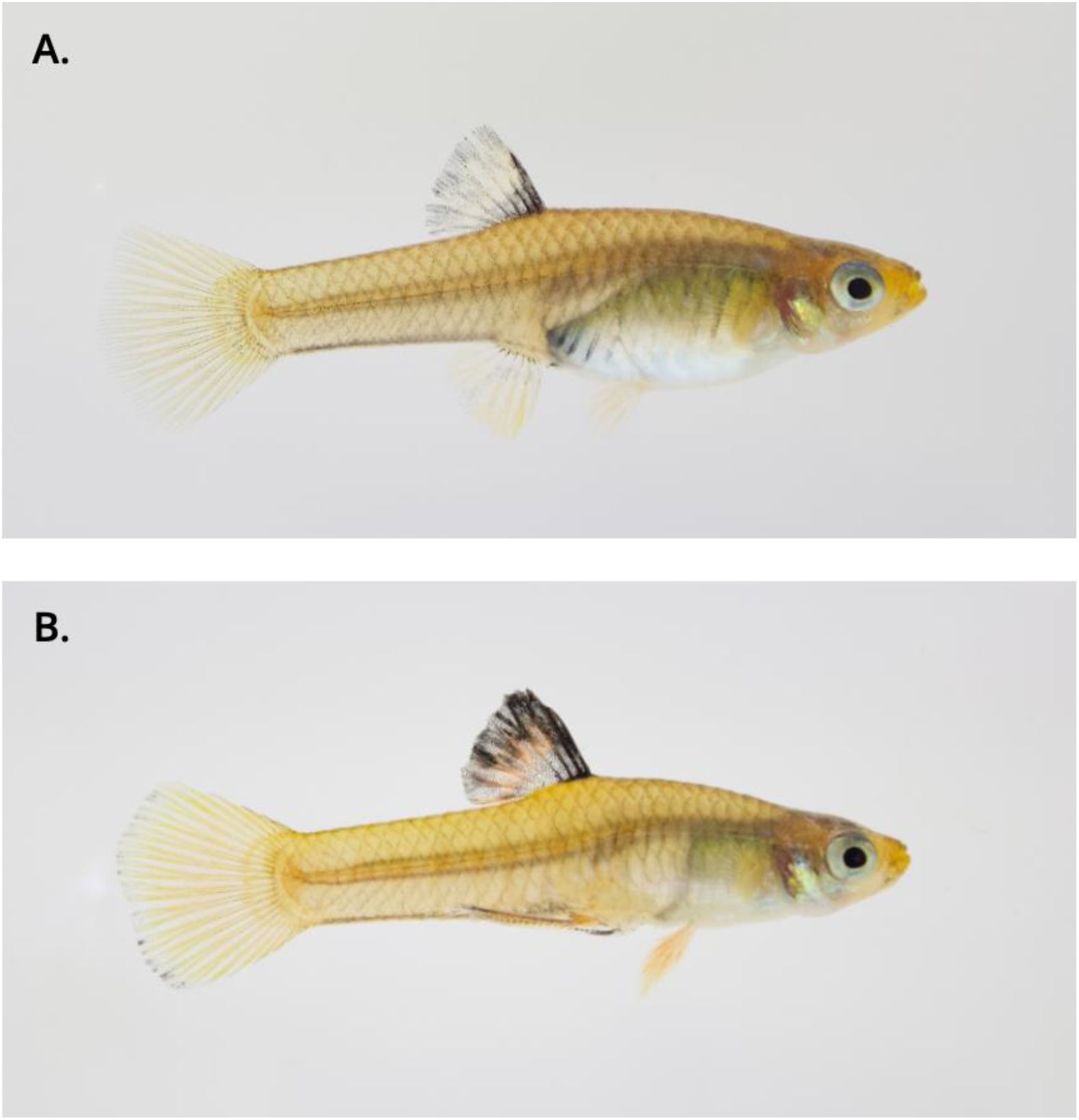
Photograph depicting mature *Xenophallus umbratilu*s (A) female and (B) male.

Specifically, in this study we address two main questions. (1) What is the life history of *Xenophallus*? To date, no studies have described any aspect of life history phenotypes in this species. We take a broad approach by describing life history traits in multiple populations throughout the geographic range of the species. (2) Are there differences in life history traits among the four phylogeographically distinct clades previously defined in *Xenophallus*? We ask if variation in life history traits exists among populations that is coincident with historical isolation and population structure. Remarkably, we find that life history traits are highly conserved among populations, despite the long periods of isolation among clades dating as far back as the Pliocene.

## Materials and methods

### Study system

Here, we focus on the livebearing fish *Xenophallus umbratilis. Xenophallus* is a monotypic species that occurs in southern Nicaragua and northern Costa Rica. It typically inhabits small headwater streams (Johnson *et al*., 2020) but has rarely also been observed in large rivers and even in Lake Nicaragua (Bussing, 1987). Previous work by Jones and Johnson (2009) using mitochondrial DNA showed that this species is divided into four distinct phylogenetic clades, with the deepest split occurring roughly 4.5 million years ago. The pattern of historical population structuring in this species appears to be influenced by bouts of marine incursion events due to historical changes in sea level over time—that is, the divergence events among clades is tied to instances where sea level rise isolated *Xenophallus* populations from each other, forcing these fishes to head water habitats in relatively large drainage basins (Jones & Johnson, 2009). Thus, populations in different clades have a long history of isolation from each other, with the longest period of isolation found between what we hereafter refer to as Clade 1 and Clades 2, 3, and 4 (Figure 2).

**Figure 2.**
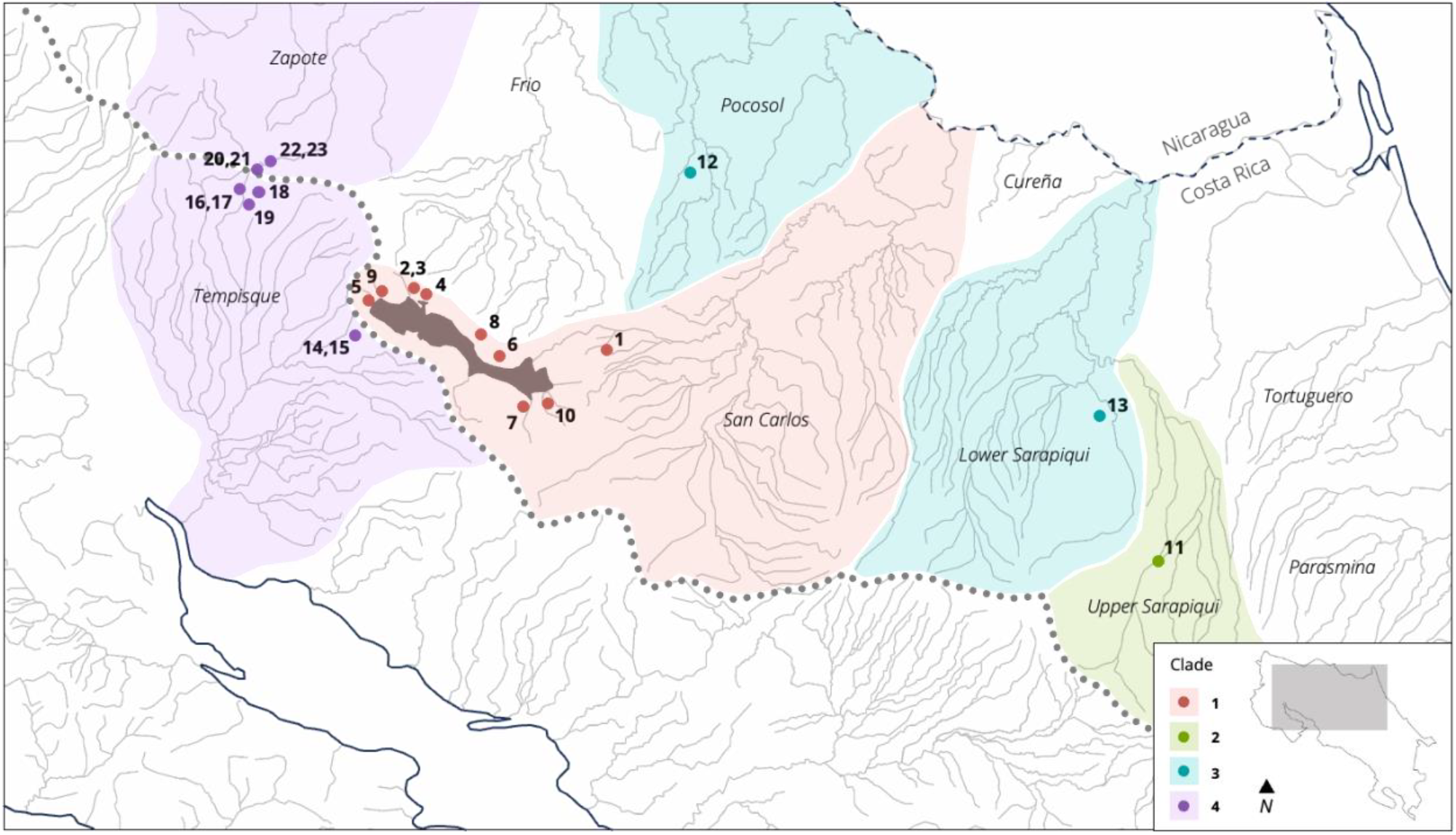
Map showing hydrological drainages in northwestern Costa Rica. Drainages are labeled by name. Drainages where life history samples were collected are shaded with colors representing phylogenetic clades as reported in Johnson and Jones (2009): red for Clade 1, green for Clade 2, blue for Clade 3, and purple for Clade 4. Numbered dots indicate populations of *Xenophallus* within these drainages where samples were collected, and population numbers correspond to those listed in Table 1. The dotted line represents the continental divide and grey lines represent major rivers.

Beyond this phylogeographic history, little is known about the basic biology of *Xenophallus*. As with other livebearing fishes (Poeciliidae), males possess a modified anal fin (gonopodium) that they use to inseminate females internally, and females carry their developing young internally until they give birth to free-swimming, juvenile offspring. The male gonopodium in this species is unusual among livebearers for being anti-symmetrical, where it terminates with a corkscrew-like twist in either a dextral or sinistral orientation (Johnson *et al*., 2020). Females have indeterminate growth, and consequently as adults are on average larger than males, which cease growing upon maturation (Meffe & Snelson, 1989). *Xenophallus* are most abundant near the headwaters of streams in habitats with relatively low flow rates, high canopy cover, and mostly devoid of piscivorous predators (Bussing, 1987), although they can occur at lower elevations where they are uncommon and less abundant (Angulo et al., 2014). Individuals appear to breed year-round. Beyond this, we know nothing about the life history traits of *Xenophallus*.

**Table 1.** Summary data from 23 *Xenophallus umbratilis* life history collections. Location information includes clade, collection ID, drainage, site name, year of collection, global positioning system (GPS) coordinates, elevation (m), and predation environment (low or high). Sample and life history data includes sample size (number of mature females and total number of individuals dissected), size class in which the majority of females were mature, brood mass, number of offspring, offspring size, and the slope of embryo mass plotted against developmental stage. Values for size at maturity, brood mass, number of offspring, and offspring are adjusted least squares means from the linear models (see methods). Negative slope values of embryo mass plotted against developmental stage indicate that this is a lecithotrophic species.

| Clade | Site ID | Drainage | Site Name | Year | GPS | Elevation | Predation | n Mature / n | Size at Maturity (mm) | Reproductive Allotment (g) | Fecundity | Offspring Size (g) | Offspring Size (mg) / Stage |
| --- | --- | --- | --- | --- | --- | --- | --- | --- | --- | --- | --- | --- | --- |
| 1 | 1 | San Carlos | Quebrada Chorros | 2024 | 10.476667°, -84.662500° | 318 | low | 15 / 24 | 24 - 26 | 0.0171 | 18.06 | 0.00114 | -0.0068 |
| 1 | 2 | San Carlos | Quebrada La Palma 1 | 2024 | 10.560808°, -84.940264° | 598 | low | 17 / 24 | 26 - 28 | 0.0101 | 12.06 | 0.00106 | -0.0043 |
| 1 | 3 | San Carlos | Quebrada La Palma 2 | 2006 | 10.560808°, -84.940264° | 598 | low | 34 / 50 | 30 - 32 | 0.0092 | 9.42 | 0.00107 | -0.0644 |
| 1 | 4 | San Carlos | Quebrada La Palmita | 2007 | 10.563400°, -84.937367° | 596 | low | 20 / 25 | 32 - 34 | 0.0089 | 13.89 | 0.00091 | -0.0565 |
| 1 | 5 | San Carlos | Quebrada Naranjos Agrios | 1998 | 10.541184°, -84.982986° | 563 | low | 27 / 38 | 32 - 34 | 0.0097 | 16.23 | 0.00087 | -0.0239 |
| 1 | 6 | San Carlos | Quebrada Perez | 2006 | 10.473611°, -84.822222° | 572 | low | 17 / 36 | 30 - 32 | 0.0191 | 25.94 | 0.00094 | -0.0749 |
| 1 | 7 | San Carlos | Quebrada Zanja | 2024 | 10.421472°, -84.769545° | 575 | low | 16 / 25 | 22 - 24 | 0.0198 | 14.91 | 0.00149 | -0.1080 |
| 1 | 8 | San Carlos | Rio Isabel | 2006 | 10.506389°, -84.845833° | 569 | high | 35 / 47 | 26 - 28 | 0.0138 | 16.12 | 0.00091 | -0.0849 |
| 1 | 9 | San Carlos | Rio Sabalito | 2006 | 10.548611°, -84.980833° | 562 | high | 39 / 50 | 26 - 28 | 0.0187 | 21.19 | 0.00091 | -0.0684 |
| 1 | 10 | San Carlos | Vuelta del Barracho | 2024 | 10.427222°, -84.752222° | 569 | low | 18 / 19 | 24 - 26 | 0.0273 | 22.79 | 0.00137 | -0.0319 |
| **Clade 1** |  |  |  |  |  |  |  |  | **27.2 - 29.2** | **0.0149** | **17.32** | **0.00105** | **-0.0524** |
| 2 | 11 | Upper Sarapiquí | Rio Corinto | 2006 | 10.211111°, -83.885278° | 231 | low | 13 / 30 | 28 - 30 | 0.0203 | 18.45 | 0.00115 | -0.0554 |
| **Clade 2** |  |  |  |  |  |  |  |  | **28 - 30** | **0.0218** | **19.75** | **0.00115** | **-0.05.54** |
| 3 | 12 | Pocosol | Rio Chimurria | 2006 | 10.727500°, -84.558333° | 62 | high | 14 / 15 | 24 - 26 | 0.0396 | 33.56 | 0.00098 | -0.0516 |
| 3 | 13 | Lower Sarapiquí | Rio Isla Grande | 2006 | 10.393055°, -83.968055° | 51 | high | 21 / 27 | 26 - 28 | 0.0195 | 20.58 | 0.00103 | -0.0316 |
| **Clade 3** |  |  |  |  |  |  |  |  | **25 - 27** | **0.0254** | **25.06** | **0.00099** | **-0.0416** |
| 4 | 14 | Tempisque | Quebrada Azul 1 | 1998 | 10.499250°, -84.985633° | 546 | low | 15 / 21 | 30 - 32 | 0.0146 | 16.91 | 0.00109 | -0.0655 |
| 4 | 15 | Tempisque | Quebrada Azul 2 | 2006 | 10.486667°, -84.986667° | 480 | low | 29 / 35 | 28 - 30 | 0.0325 | 28.42 | 0.00126 | -0.0678 |
| 4 | 16 | Tempisque | Quebrada Hormiguero 1 | 2005 | 10.690833°, -85.083611° | 501 | low | 15 / 24 | 36 - 38 | 0.0156 | 14.99 | 0.00127 | -0.0597 |
| 4 | 17 | Tempisque | Quebrada Hormiguero 2 | 2006 | 10.690833°, -85.083611° | 501 | low | 25 / 47 | 32 - 34 | 0.0211 | 16.61 | 0.00134 | -0.0854 |
| 4 | 18 | Tempisque | Rio Esquivetto | 2006 | 10.687222°, -85.066667° | 493 | low | 29 / 50 | 28 - 30 | 0.0149 | 11.62 | 0.00146 | -0.0991 |
| 4 | 19 | Tempisque | Rio Tenorio | 2006 | 10.688055°, -85.076111° | 480 | low | 36 / 43 | 26 - 28 | 0.0242 | 16.33 | 0.00158 | -0.0944 |
| 4 | 20 | Zapote | Trib. to Rio Bijagua 1 | 2005 | 10.723798°, -85.066495° | 439 | low | 16 / 23 | 26 - 28 | 0.0128 | 13.53 | 0.00119 | -0.1110 |
| 4 | 21 | Zapote | Trib. to Rio Bijagua 2 | 2006 | 10.724167°, -85.066389° | 439 | low | 39 / 40 | 24 - 26 | 0.0228 | 14.41 | 0.00179 | -0.1060 |
| 4 | 22 | Zapote | Trib. to South Rio Bijagua 1 | 2006 | 10.731199°, -85.055627° | 452 | low | 16 / 21 | 24 - 26 | 0.0133 | 12.11 | 0.00137 | -0.1190 |
| 4 | 23 | Zapote | Trib. to South Rio Bijagua 2 | 2006 | 10.731199°, -85.055627° | 452 | low | 42 / 49 | 26 - 28 | 0.0179 | 13.84 | 0.00141 | -0.0914 |
| **Clade 4** |  |  |  |  |  |  |  |  | **28 - 30** | **0.0197** | **16.86** | **0.00137** | **-0.0899** |

### Sampling

To test for life history diversification within *Xenophallus*, we collected 23 samples across 19 sites throughout the range of this species (Figure 2). Within Clade 1, we sampled fish from the San Carlos drainage (which includes tributaries to Lake Arenal), and within Clade 4, we sampled fish from the Zapote and Tempisque drainage. We also sampled fish from three localities within the Sarapiqui and Pocosol drainages, which include representatives of Clades 2 and 3; populations in these clades are not common, resulting in fewer sites than found in Clades 1 and 4 (see Table 1). We nevertheless included these samples to see how life history traits from these populations compared to those from the more broadly represented clades.

In most, but not all cases, we sampled locations within the drainages near headwater habitats found along the volcanic cordillera in northern Costa Rica. Previous work in this region has shown that piscivorous predators are typically absent from this type of habitat, resulting in what has sometimes been called a ‘low predation’ environment (Reznick & Endler, 1982; Johnson & Belk, 2001; Johnson *et al*., 2002). This absence of piscivorous predators typically results in a reduced mortality schedule relative to locations where predators are present (Reznick *et al*., 1996; Johnson & Zuñiga-Vega, 2009). Because the piscivorous predation regime is known to impact life history, we scored each population as high or low predation risk based on the presence of the common piscivorous predator, *Parachromis dovii*, and included this as a factor in our models. Additionally, we sampled all sites for this study during the dry season (January to April) to avoid potential plastic phenotypic shifts in life histories due to differences in resource availability associated with changes in resource abundance (Johnson *et al*. 2024).

We collected 23 samples of *Xenophallus*, 10 from nine sites representing Clade 1, one from a site representing Clade 2, two from two sites representing Clade 3, and 10 from six sites representing Clade 4. We collected up to 100 fish at each site using handheld seines. We euthanized fish in the field using MS-222 (a fish anesthesia) and immediately preserved individuals in 70% ethanol. We haphazardly collected individuals across the full-size distribution of males, females, and juveniles, to represent all size classes. We transported these preserved specimens to the Evolutionary Ecology Laboratories at Brigham Young University where all samples were processed. All sampling was conducted following guidelines approved by the BYU Institutional Animal Care and Use Committee (IACUC #21-1118) and with permission from the Costa Rica Ministrio del Ambiente y Energía (MINAE), Sistema Nacional de Áreas de Conservación (SINAC).

### Data collection

We measured life history traits in each of the populations following the protocol of Johnson (2025). For each female, we measured standard length, reproductive allotment, fecundity, and offspring size. To collect these data, we divided females into 2-mm size classes. To calculate size at maturity, we identified the size class in which at least half of the individuals contained developing embryos (defined here as stage 4 or greater following Haynes (1995)). We calculated reproductive allotment as the dry mass of a single brood of offspring and fecundity as the total number of individuals in a developing brood. We calculated offspring size as the average per-capita dry mass of developing offspring. Only females with developing embryos were included in the estimates of reproductive allotment, number of offspring, and offspring size. We measured dry mass for both embryos and adult females (with embryos and digestive tract removed) after 24 h in a 55°C desiccating oven.

### Data analysis

We used linear mixed models to evaluate whether differences in life history trait expression exist between the four clades using the lmer() function from the lme4 package in R (Bates *et al*., 2015). This package uses Satterthwaite’s method to calculate effective degrees of freedom, which can result in partial degrees of freedom (Bates *et al*., 2015). In all tests, we used an alpha value of 0.05. All analyses were conducted in R v.4.2.2 (R Core Team 2020). Prior to analysis, we transformed raw data for each of the following traits as follows (transformations in parentheses): female dry mass (log_10_), female standard length (log_10_), brood dry mass (log_10_), and offspring count (square-root). For fecundity, we included clade, standard length, and predation environment as fixed effects and site as a random effect. For reproductive allotment and embryo size, we included standard length, predation environment, and embryo stage as fixed effects and site as a random effect. For female size at maturity, and for the slope of embryo mass across developmental stages, we used a linear model with clade as a fixed effect. In linear models where ‘clade’ was significant, we conducted a Tukey’s post hoc test to determine which of the four clades differed from each other.

## Results

We tested if populations of *Xenophallus* from four evolutionary clades differed for any life history traits: they mostly did not. There were no differences in female life history traits of size at maturity, reproductive investment, and number of offspring among any of the four clades (Figure 3; Table 1). The only differences we found were that offspring size was larger in Clade 4 than in Clade 1, and that the rate of decrease in embryo mass throughout embryonic development (measured by slope) was greater in Clade 4 than in Clade 1 (Figure 3). Here, we present the results of the linear models evaluating each trait. We also summarize these data, providing the first ever life history description of this species (Table 1).

**Figure 3.**
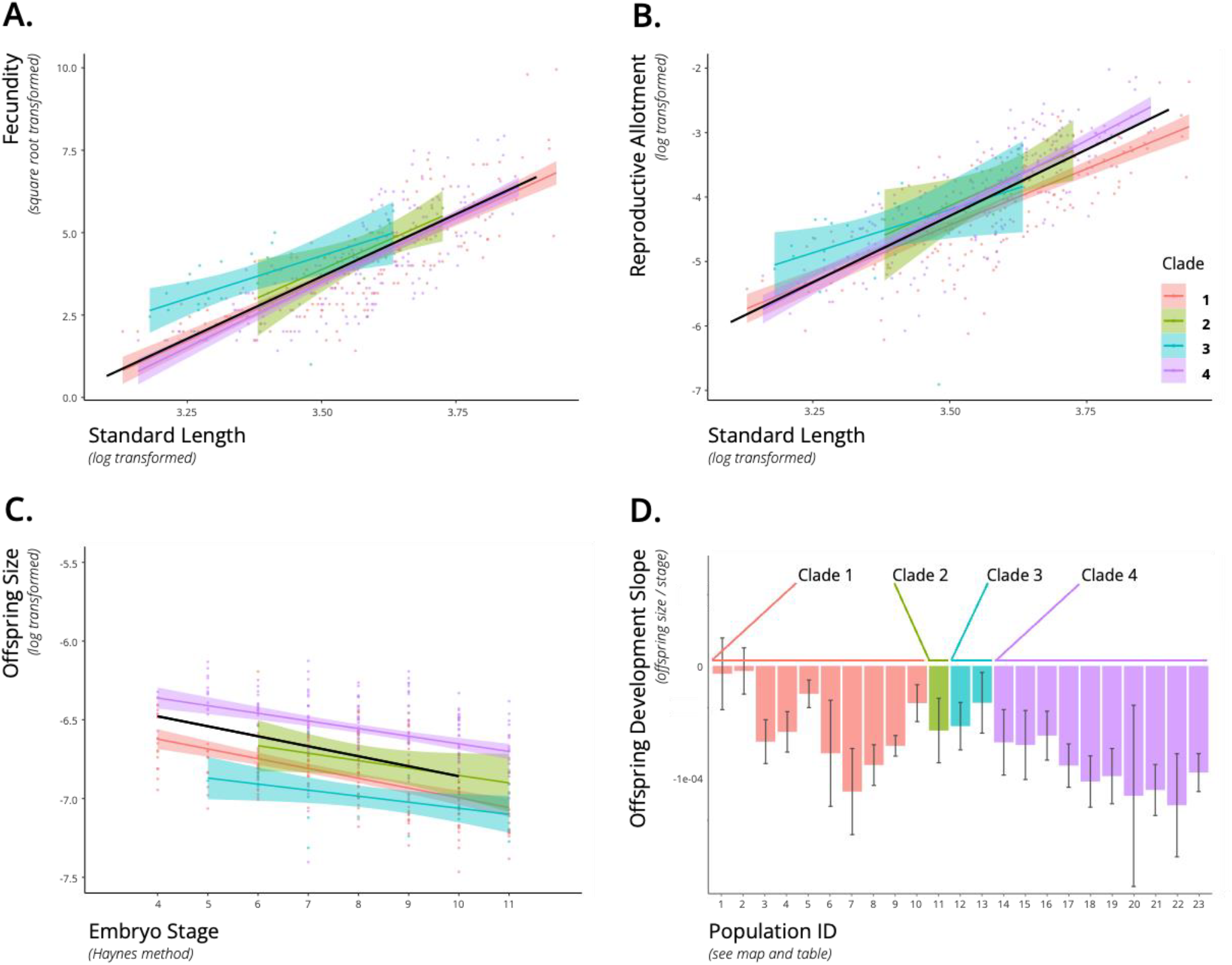
*Xenophallus* life history traits showing raw data for individual females for fecundity, reproductive allotment, embryo mass, and embryo mass across development. **A. Fecundity:** Plot showing relationship between fecundity (sqrt transformed) and female standard length (log transformed) for individuals from Clades 1, 2, 3, and 4 (identified by color; see inset key). **B. Reproductive allotment:** Plot showing relationship between female reproductive allotment (log transformed) and female standard length (log transformed) for individuals from Clades 1, 2, 3, and 4. **C. Embryo mass:** Plot showing relationship between embryo mass (log transformed) and embryo developmental stage (Haynes method) for embryos extracted from individuals from Clades 1, 2, 3, and 4. **D. Embryo mass across development:** A plot showing the change in embryo mass across developmental stages (slopes) for embryos by population coded by clades (see Figure 2 for clades).

### Life history traits

There were no significant differences among clades for reproductive allotment ( x_clade1_ = 0.0149g, x_clade2_ = 0.0219g, x_clade3_ = 0.0254g, x_clade4_ = 0.0197g; Tukey’s post hoc for all clade comparisons, p > 0.05), fecundity ( x_clade1_ = 17.32, x_clade2_ = 19.76, x_clade3_ = 25.06, x_clade4_ = 16.86; Tukey’s post hoc for all clade comparisons, p > 0.05), and female size at maturity ( x_clade1_ = 27.2-29.2mm, x_clade2_ = 28-30mm, x_clade3_ = 25-27mm, x_clade4_ = 28-30mm; p = 0.746). There was a significant difference between Clades 1 and 4 in embryo size, with Clade 1 having smaller embryos than Clade 4 (x_clade1_ = 0.00105g, x_clade2_ = 0.00115g, x_clade3_ = 0.00101g, x_clade4_ = 0.00137g, Tukey’s post hoc: Clade 1 vs. Clade 4 comparison p = 0.0081; for all other clade comparisons, p > 0.05). *Xenophallus* does not express superfetation—that is, females only develop one single brood at a time. We noted that embryo mass decreased across developmental stages in all populations tested from all clades, indicating a lecithotrophic maternal resource allocation strategy. We also found a significant difference between the rate of decrease in embryo mass between populations from Clade 1 and Clade 4 ( x_clade1_ slope = -0.0664, x_clade2_ slope = -0.0554, x_clade3_ slope = -0.0365mg / stage, x_clade4_ slope = -0.0703; Tukey’s post hoc: clade1 vs.clade 4 comparison p = 0.0317; for all other clade comparisons, p > 0.05).

## Discussion

### Conservation units and assessment

A key challenge in conserving standing biodiversity is knowing how to delineate populations within species that might warrant unique management or protection (Shafer *et al*. 2015). Conservation biologists have used the concept of ‘conservation units’ to frame these decisions, and there is a rich body of work exploring how to define and utilize conservation units under various laws or management frameworks (reviewed in Hoelzel (2023); e.g., United States’ Endangered Species Act (U.S. Congress 1973) and IUCN Red List (IUCN Standards and Petitions 2024)). One common approach is to use phylogenetic data to delineate groups of populations that share distinct evolutionary histories, which can be viewed as hypothetical conservation units (Crandall *et al*. 2000). Such units can then be tested using data from functional traits to evaluate their validity. Life history traits are useful in this regard as they are known to evolve rapidly in response to a variety of selective pressures (Gallagher *et al*., 2021). In fact, life histories are often the first traits to diverge among distinct evolutionary units and therefore provide the first independent evidence to support targeted conservation and management actions.

We found that life history traits in *Xenophallus* showed very little divergence among the four clades considered here, even among clades isolated for 4.5 million years. We found only modest differences in life history traits between Clade 1 and Clade 4 (Figure 3). Females from Clade 1 gave birth to offspring that were slightly larger in size than those in Clade 4; and developing embryos from Clade 4 decreased in size more rapidly than those in Clade 1. Hence, these two clades show modest divergence from each other. Yet, neither of these clades differed from Clades 2 or 3; and no other life history traits differed among any of the clades. It is difficult to draw conclusions about Clades 2 and 3, as we only sampled a few populations due to their rarity (see Angulo *et al*., 2013); however, the populations that we did sample fell directly within the range of traits from other populations from clades 1 and 4, suggesting that these clades may not be evolutionarily distinct; additional data from populations in these lowland drainages (Figure 2) are be needed to further explore this conclusion.

Although the documented range of this species extends from small-order streams at high elevations (∼500 m above sea level) to larger rivers at low elevation (around 50 m above sea level) (Bussing, 2002), our sampling suggests that *Xenophallus* is common and most abundant at higher elevations, and uncommon or absent at lower elevations. The most recent conservation assessment suggesting that populations of *Xenophallus* are exceptionally rare (<10 populations) (Lyons and McMahan, 2019) is clearly not correct. In our study alone, we sampled *Xenophallus* from 19 localities, but primarily from high-elevation locations (Figure 2; Table 1). Moreover, individuals are abundant in these high-elevation habitats, which coincidentally have little human impact (pers. obs.). In contrast, *Xenophallus* is absent or uncommon at low-elevation habitats, which tend to be more impacted by agriculture and development (Angulo et al., 2013; Lyons and McMahan, 2019). Consequently, it is not clear if the relative rarity of *Xenophallus* at lower elevations is due to anthropogenic causes, or if it is a consequence of natural phenomenon, including historical marine incursion events over evolutionary time (Jones & Johnson, 2009) and more complex ecological interactions with competitors and predators at these sites. Whatever the case, *Xenophallus* appears to be uncommon at lower elevation habitats, which happen to mostly comprise Clades 2 and 3 (Figure 2).

### Life history of Xenophallus umbratilis

Ours is the first study to describe the life history of *Xenophallus*. This species shows a life history typical of other species in the Poeciliid family (Furness *et al*., 2019). *Xenophallus* is lecithotrophic; that is, females fully provision eggs prior to fertilization with no additional resources to developing embryos post fertilization (Johnson and Bagley, 2011). This species also does not exhibit superfetation, which is the ability of females to simultaneously carry multiple broods of developing young, each at different stages of development (Johnson and Bagley, 2011). Instead, *Xenophallus* carries only one developing brood at a time. Previous work suggests that lecithotrophy and non-superfetation is the ancestral state of poeciliid fishes (Furness *et al*., 2019). Other life-history traits such as fecundity, size at maturity, and reproductive allotment in *Xenophallus* are also typical in comparison to life-histories of other poecilids (Johnson & Bagley, 2011).

What is important to note in the life history of *Xenophallus* with respect to conservation is that females mature relatively early and give birth to large broods of offspring. Females also reproduce year round. This high level of fecundity typically means that population growth can be achieved fairly quickly. To specifically calculate population growth rates in *Xenophallus* will require age-specific estimates of mortality rates (*sensu* Johnson & Zuñiga-Vega, 2009) combined with the life history data that we have presented here. Estimating mortality rates can be achieved in different ways, including serial mark-recapture experiments (Dorazio & Rago, 1991; Colchero et al., 2012). Previous demographic work in other livebearing fishes suggests that in the absence of predators, populations typically show positive growth rates (Reznick & Bryga, 1996; Johnson & Zuñiga-Vega, 2009). This suggests that *Xenophallus* from high-elevation localities that lack fish predators (see Table 1) are likely sustainable under current conditions. Although population sizes are robust in these locations, protecting these habitats should be a high priority for the long-term persistence of *Xenophallus*.

### Conclusions and future direction

Knowing the life history of *Xenophallus* advances our ability to conserve this species in important ways. We found that although *Xenophallus* is composed of at least four phylogenetically distinct clades, life history traits are mostly conserved in this species. Two clades with populations that occupy high-elevation habitats show some life history differences. To better understand the broad conservation status of *Xenophallus* across its range will require more exhaustive sampling in drainages that comprise low-elevation habitats, including the lower reaches of the San Carlos, the Pocosol, and the Sarapiqui drainages (Figure 2). Estimating mortality rates in these populations, as well as in populations in high-elevation habitats, will allow us to determine the long-term stability of populations, as well as their potential for growth. *Xenophallus* is one of many Central American freshwater fishes that comprise the fish fauna of this region. Continued efforts to document the distribution and abundance of these fishes, their patterns of historical and contemporary gene flow, and demographic parameters (including life history traits) present the next steps to understanding and conserving the remarkably diverse freshwater fishes of the Neotropics.

## Acknowledgements

We appreciate the support of Javier Guevara Siquiera and Lourdes Vargas Fallas at Vida Silvestre, Ministerio del Ambiente y Energía (MINAE), Sistema Nacional de Áreas de Conservación (SINAC), Costa Rica, for processing collecting permits for most of the specimens used in this study. We are also grateful to the field researchers who collected samples that were made available for our use from the BYU Life Science Museum, including Mark Belk and Jerry Johnson. We extend our gratitude to all the undergraduates at Brigham Young University and the University of Missouri–St. Louis who contributed to this research in both field and laboratory work. E. Elias Johnson received funding from the College of Life Sciences, Brigham Young University and Erik S. Johnson was funded in part by the Whitney R. Harris World Ecology Center.

